## Supplementary material for "Glial plasticity and metabolic stability after knockdown of astrocytic Cx43 in the dorsal vagal complex": Table S1

| Gène | | Primer Sequence | Target Site | qPCR T° | Amplicon Size | mRNA Identifier |
|---|---|---|---|---|---|---|
| **ywhaz** | F | ACATCTGCAACGATGTACTGTCT | 680-702 (Exon 3-4) | 60.0°C | 166 bp | NM_001253805 |
| | R | TGCTGTGACTGGTCCACAAT | 826-845 (Exon 4) | | | |
| **mGjb2** (Cx26) | F | AGCCGTCTTCATGTACGTCTTT | 697-718 (Exon 2) | 60.0°C | 124 bp | NM_008125.3 |
| | R | CTTTTCTGTGGGCCTGGAAATG | 799-820 (Exon 2) | | | |
| **mGjb6** (Cx30) | F | GGCCAACTGAGAAAACGGTG | 1247-1266 (Exon 3) | 60.0°C | 56 bp | NM_008125.3 |
| | R | GCAAATCACGGATGCGGAAA | 1283-1302 (Exon 3) | | | P000002.166.f ou r |
| **Gjb1 (cx32)** | F | GTGGACCTATGTCATCAGTGTGG | 449-471 (exon 2) | 60.0°C | 119 bp | NM_001302496 |
| | R | GGAAGGCTTCACACTTGACCAG | 546-567 (exon 2) | | | |
| **Cx36** | F | TGATTGGGAGGATCCTGTTGAC | 547-568 (exon 1) | 60.0°C | 95 bp | BC058595 |
| | R | CATGGTCTGCTCATCATCGTAC | 620-641 (exon 1) | | | |
| **mGja1** (Cx43) | F | GGTGGACTGCTTCCTCTCAC | 817-836 (Exon 2) | 60.0°C | 151 bp | NM_010288.3 |
| | R | ATCGCTTCTTCCCTTCACGC | 948-967 (Exon 2) | | | |
| **Gjc1** (Cx45) | F | TTGGGTAACAGGAGTTCTGGTGAA | 789-812 (Exon 2) | 60.0°C | 145 bp | NM_008122.2 |
| | R | GTGAGCCAGATCTTCCCTACA | 892-912 (Exon 3) | | | |
| Gjc2 (Cx47) | F | GAGGATGAGGACGAGGAACCA | 1275-1295 (exon 1) | 60.0°C | 111 bp | XM_036156249.1 |
| | R | CACCGTCTTTCCATCACCTCC | 1365-1385 (exon 1) | | | |
| **Panx1** | F | CAGGCTGCCTTTGTGGATTC | 667-686 (Exon 2) | 60.0°C | 145 bp | NM_019482 |
| | R | CGGGCAGGTACAGGAGTATG | 792-811 (Exon 3) | | | |
| **GLT1 (Slc1a2)** | F | CCAACAATATGCCCAAGCAGG | 616-636 (exon2) | 60.0°C | 155 bp | NM_001077514 |
| | R | TGCTCCCAGGATGACACCAA | 751-770 (exon 2-3) | | | |
| **GLAST (Slc1a3)** | F | CACTGCTGTCATTGTGGGTA | 731-750 (exon 2-3) | 60.0°C | 125 bp | NM_ 600111 |
| | R | CCATTCCTGTGACGAGACT | 873-891 (exon 3-4) | | | |
| **mGluR1 (Grm1)** | F | AGGGCGATGCTTGATATCGT | 1038-1057 (exon2) | 60.0°C | 88 bp | NM_001114333 |
| | R | CCATTCCACTCTCGCCGTAA | 1106-1125 (exon3) | | | |
| **mGluR2 (Grm2)** | F | GCGGCTCCTACAGTGATGTC | 641-661 (exon 2) | 60.0°C | 134 bp | NM_001160353 |
| | R | TCATAACGGGACTTGTCGCTC | 756-775 (exon3) | | | |
| **mGluR3 (Grm3)** | F | ACCAAGCTCTGTGATGCAATG | 2245-2265 (exon4) | 60.0°C | 142 bp | NM_001417964 |
| | R | TCCCGTCTCCGTAAGTGTCA | 2367-2386(exon5) | | | |
| **mGluR4 (Grm4)** | F | GAAGGTGCAGTCACCATTCTTC | 1959-1980 (exon5) | 60.0°C | 101 bp | NM_001291045 |
| | R | AACCAGATGTTGCGCCTGTT | 2040-2059 (exon6) | | | |
| **mGluR5 (Grm5)** | F | CCCGAGCCATGGTAGACATA | 1311-1330 (exon3) | 60.0°C | 140 bp | NM_001143834 |
| | R | AGAGTGGGCGATGCAAATCC | 1431-1450 (exon4) | | | |
| **mGluR6 (Grm6)** | F | CCTGCTGTTGGCACTGTGA | 1752-1770 (exon 9) | 60.0°C | 114 bp | NM_173372 |
| | R | GACGGCAGCCAGTGTGGTT | 1752-1770 (exon 9) | | | |
| **mGluR7 (Grm7)** | F | GGCATTGGGCTGGAATTATGT | 912-932 (exon 2) | 60.0°C | 127 bp | NM_177328 |
| | R | AATTCTCACGGACTGGGCAAT | 1018-1038 (exon 3) | | | |
| **mGluR8 (Grm8)** | F | TACAGCTCCTtGGAGTGTT | 2546-2564 (exon 9) | 60.0°C | 71 bp | NM_001361125 |
| | R | CTGTTCtCCATAGTCAATGAT | 2596-2616 (exon 9) | | | |

**Table S1:** Primers sequences used for SYBR Green assays.
