## Supplementary material for "Glial plasticity and metabolic stability after knockdown of astrocytic Cx43 in the dorsal vagal complex": Figure S1

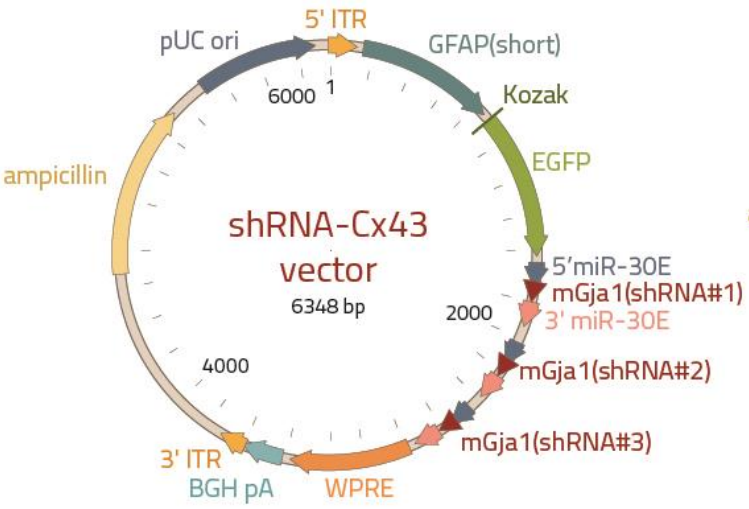


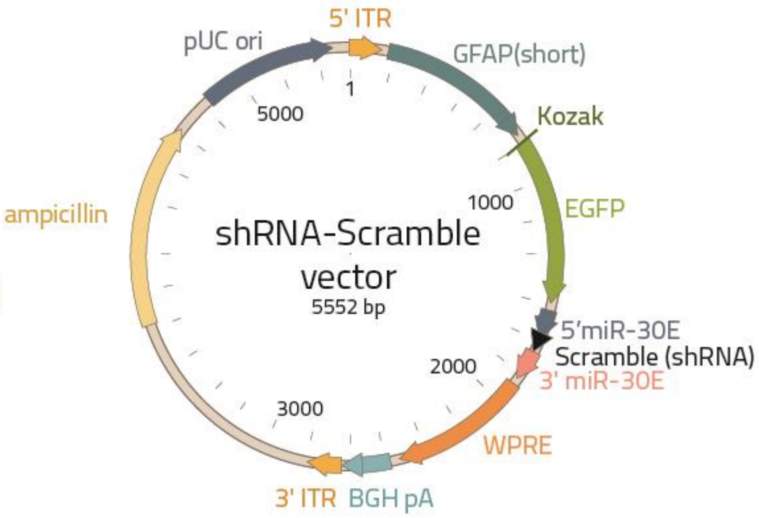


| | **ShRNA-Cx43** | **ShRNA-Scramble** |
|---|---|---|
| Vector name | pAAV[3miR30]-GFAP(short)>EGFP:{mGja1[shRNA#1]}:{mGja1[shRNA#3]}:{mGja1_shRNA}:WPRE | pAAV[miR30]-GFAP(short)>EGFP:Scramble[miR30-shRNA#1]:WPRE |
| Vector size | 6348 bp | 5552 bp |
| Target sequences by shRNA | GCCTGATGACCTGGAGATTTAA<br>ACAATTCCTCCTGCCGCAATTA<br>GAACAGTCTGCCTTTCGCTGTA | ACCTAAGGTTAAGTCGCCCTCG |
